# Spontaneous Mutation in *2310061I04Rik* Results in Reduced Expression of Mitochondrial Genes and Impaired Brain Myelination

**DOI:** 10.1101/2023.08.10.552737

**Authors:** Erdyni N. Tsitsikov, Khanh P. Phan, Yufeng Liu, Alla V. Tsytsykova, Rosalia Paterno, David M. Sherry, Anthony C. Johnson, Ian F. Dunn

## Abstract

Here, we describe a spontaneous mouse mutant with a deletion in a predicted gene *2310061I04Rik* (*Rik*) of unknown function located on chromosome 17. A 59 base pair long deletion occurred in the first intron of the *Rik* gene and disrupted its expression. *Rik^null^* mice were born healthy and appeared anatomically normal up to two weeks of age. After that, these mice showed inhibited growth, ataxic gait, and died shortly after postnatal day 24 (P24). Transcriptome analysis at P14 and P23 revealed significantly reduced expression of mitochondrial genes in *Rik^null^* brains compared to wild type controls including *mt-Nd4*, *mt-Cytb*, *mt-Nd2*, *mt-Co1*, *mt-Atp6,* and others. Similarly, genes specific for myelinating oligodendrocytes also showed reduced expression in P23 *Rik^null^* brains compared to controls. Histological examination of anterior thalamic nuclei demonstrated decreased myelination of anteroventral nuclei but not of anterodorsal nuclei in P23 *Rik^null^* mice. Myelination of the anterior commissure was also impaired and displayed extensive vacuolation. Consistent with these findings, immunohistochemistry showed reduced expression of Opalin, a glycoprotein expressed in differentiated oligodendrocytes. Taken together, these results suggest that RIK is important for oligodendrocyte maturation, and myelination in the developing brain.

## Introduction

Neuroscience has been greatly advanced by the availability of mutant mouse models. For many years, various spontaneous mutations and polymorphic alleles provided the only source for genetic studies. Classic neurological mouse mutants include *shiverer* (Roach et al., 1983), *quaking* (Ebersole et al., 1996), *reeler* (Hartfuss et al., 2003), *pcd* (Mullen et al., 1976) and many others, which have been critical tools for understanding the function of specific genes in the central nervous system. Affected homozygotes of *shiverer* (*shi*) mice develop characteristic “shaking” or “shivering” gait approximately two weeks after birth. This shivering increases in severity with age, and mice die prematurely, typically between 50 and 100 days after birth (Readhead and Hood, 1990). The central nervous system (CNS) of the mutant mouse is hypomyelinated but the peripheral nervous system (PNS) appears normal. These mice fail to make myelin due to a deletion of the gene, encoding myelin basic protein (MBP) (Readhead and Hood, 1990).

Homozygous *quaking viable* (*qk^v^*) mice display vigorous tremors starting at about postnatal day 10 (P10), especially pronounced in hind limbs, and experience convulsive tonic-clonic seizures as they mature (Haroutunian et al., 2006, Ebersole et al., 1996). These mice have pronounced demyelination in both CNS and PNS, resulting from reduced numbers of myelin lamellae and failure of the resulting myelin to compact properly (Suzuki and Zagoren, 1977). The original *qk^v^* allele resulted from a large spontaneous deletion that affects three loci on chromosome 17 (chr17): *parkin* (*Prkn*), *Park2*-coregulated gene (*Pacrg*), and KH domain RNA binding quaking homolog (*Qki*), which is highly expressed in oligodendrocytes (OLs) and astrocytes in CNS as well as Schwann cells in PNS (Hardy et al., 1996). Further analysis revealed that the alterations in *Qki* locus are responsible for neurological phenotype of *qk^v^* mice (Noveroske et al., 2005). Forward genetics experiments generated additional *Qki* locus mutants with varying phenotypes depending on genetic environment of any given allele. Those phenotypes are also compounded by complex multigenic differences among commonly used mouse strains and substrains (Yoshiki and Moriwaki, 2006, Mekada and Yoshiki, 2021).

In this study, we describe a serendipitous mutation in a predicted gene, *2310061I04Rik*, which causes a similar neurological phenotype. *Rik* has no previously described function and is positioned on chr17 just ∼25MB downstream from the *Qki* locus. A 59-bp deletion in the first intron of the *Rik* gene spontaneously arose in one of the embryonic cell (ES) clones during generation of *Traf7^fl/fl^* mice and resulted in *Rik* expression deficiency. *Rik^null^* mice were born healthy, but at two weeks of age started to look smaller than their littermates, exhibited ataxic gait, and died before 4 weeks of age. While P14 *Rik^null^* mice demonstrated normal expression of genes specific for differentiated oligodendrocytes, they had lower expression of mitochondrial genes in compared to wild type controls. At P23, *Rik^null^* mice these mice displayed decreased expression of mitochondrial genes as well as myelinating oligodendrocyte genes. Histological and immunohistochemical studies revealed reduced myelination of anteroventral nuclei of thalami and of anterior commissure in *Rik^null^* brains.

## Results

### Spontaneous deletion in *2310061I04Rik* gene

To investigate the physiological function of TRAF7 *in vivo*, we generated mice with a conditionally targeted *Traf7* allele (Tsitsikov et al., 2023). The conditional allele contained loxP sites flanking exons 2 and 14 in the *Traf7* gene. While heterozygous *Traf7*-floxed allele (*Traf7^+/fl^*) mice were healthy and fertile, homozygous *Traf7^fl/fl^* pups appeared normal at birth, but unexpectedly stopped gaining weight after postnatal day 12 (Figure S1A and S1B) and developed a pronounced ataxia with seizure-like behavior (Movie 1) and had to be euthanized shortly after P30 (Figure S1C). For clarity, *Traf7*-floxed mice showing pathology are referred to as *Traf7^fl*/fl*^* mice to distinguish them from healthy *Traf7^fl/fl^* animals, which were described previously in Tsitsikov et al. (Tsitsikov et al., 2023). To determine whether *Traf7^fl*/fl*^* mice had floxed allele-linked spontaneous mutation(s) in protein coding regions, we performed whole exome sequencing using tail DNA from two P30 *Traf7^fl*/fl*^* animals that showed pathology and two C57BL/6 mice of the same age. None of *Traf7^fl*/fl*^* animals revealed relevant homozygous sequence variations in the coding region of genes located in the vicinity of *Traf7* on chr17 (data not shown).

Next we produced mice with whole body deletion of *Traf7* by crossing *Traf7-floxed* mice with *E2a-Cre*-trangenic animals (Lakso et al., 1996). While *Traf7^−*/+^* mice were phenotypically normal and fertile, *Traf7^−*/−*^* animals died around day 10 of embryonic development (E10) (Tsitsikov et al., 2023). To understand gene expression differences between E9.5 WT and *Traf7^−*/−*^* embryos, we compared their transcriptome profiles by RNA sequencing (RNA-seq) (Supplemental file 1). The analysis revealed that the WT embryos formed a tight cluster following principal component analysis (PCA) (Figure S2A), but the distribution of the *Traf7^−*/−*^* embryos did not cluster tightly, with three samples near the WT cluster, suggesting a recent branching of transcriptional profiles. Indeed, most identified genes (11,755) were common between these two groups (Figure S2B). Only 277 and 163 of the differentially expressed genes (DEGs) were specifically expressed in either *Traf7^−*/−*^* or WT embryos, respectively. Hierarchical clustering based on gene expression profiles (Heat map) revealed high similarity of expression patterns between samples within each group and substantial differences between them (Figure S2C). As expected, *Traf7* had the greatest significant difference in relative expression between WT embryos and their *Traf7^−*/−*^* counterparts (Figure S2D). The DEG with the largest difference in relative expression and high statistical significance was *Klf2,* a key blood flow-responsive gene in endothelial cells (Novodvorsky and Chico, 2014). Interestingly, one of the DEGs was the predicted *2310061I04Rik* gene (Figure S2D), which has an unknown function and is located on chr17 approximately 11MB downstream of *Traf7* gene (Figure 1A). The detailed analysis with the Integrative Genomics Viewer (IGV) demonstrated a 59-base pair (bp) deletion in the first intron of the *2310061I04Rik* gene, 125 bp upstream of exon 2. Curiously, this region contains recognition sequences for three commonly used restriction endonucleases Hind III, Pvu II, and Bcl I (Figure 1A). To examine the origin of this 59-bp deletion, we designed a genotyping PCR assay around this deletion and tested embryonic stem (ES) cell clones, which were used for blastocyst injection during generation of the *Traf7^fl/+^* mice. The ES124 clone was heterozygous for the *2310061I04Rik* mutation, but four other clones (ES171, ES242, ES262, and ES353) and the paternal ES cell line IN2 used for transfection, possessed no such deletion (Figure 1B, lanes 1 to 7). The *2310061I04Rik* gene was first sequenced and predicted by the RIKEN (Designated National Research and Development Institute in Japan) mouse genome encyclopedia project (Hayashizaki, 2003). Because *2310061I04Rik* has no designated name or described function, for convenience we temporarily named this gene “*Rik”,* even though it is a common suffix in names of many unknown genes identified by RIKEN.

**Figure 1.**
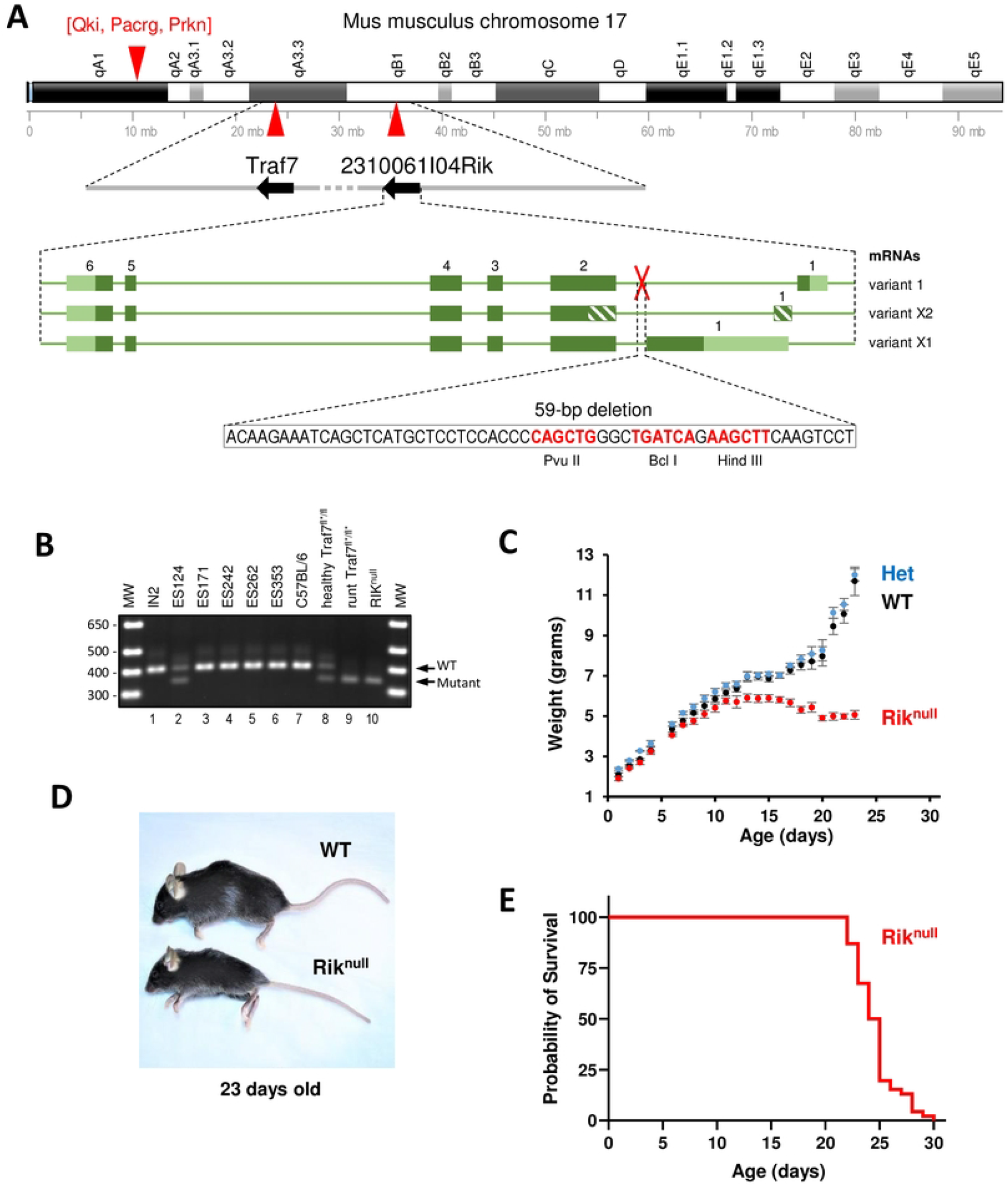
Severe runting and early death of mice with serendipitous deletion in *2310061I04Rik* gene. (**A**) Schematic drawing (up-to-scale) of the mouse chromosome 17. Location of *2310061I04Rik* and *Traf7* genes on the chromosome are marked by red triangles and their directions of transcription are shown by black arrows on grey line in a blowout drawing of their loci below. Predicted variants of mRNA transcribed from the *2310061I04Rik* gene are depicted on the bottom panel in green color. Exons presented as green boxes with coding (dark green) and non-coding (striped or light green) regions. Location of the deletion is marked by a red cross and its sequence is shown below the mRNA scheme in a blow-out window. Restriction sites within the deletion region are shown in red font with corresponding enzymes in *Italic* font underneath. (**B**) Detection of WT (401 bp) and mutant *Rik* (342 bp) alleles in ES cell line (IN2), ES clones with *Traf7*-floxed alleles, and *Traf7^fl*/fl*^* and *Rik^null^* mice by conventional PCR and agarose gel electrophoresis. MW: Molecular Weight markers (1 Kb Plus Ladder). (**C**) Mouse weight chart from birth to end of life of *Rik^null^* mice (n=175 WT, n=348 HET, n=145 *Rik^null^*, median±SEM). (**D**) Representative 23 days old WT and *Rik^null^* mice. (**E**) Probability of survival of *Rik^null^* mice (n=45).

### A 59-bp deletion in *2310061I04Rik* gene confers the runt phenotype

To explore whether the runt phenotype of *Traf7^fl*/fl*^* pups arose due to impaired expression of *Rik*, we bred *Traf7^+/fl*^* animals with C57BL/6 mice for several generations to unlink the mutated *Rik* and *LoxP3*-modified *Traf7* loci. The resulting mutant mice had a *Rik* deletion but an intact *Traf7* gene (*Rik^null^*). As shown in Figure 1B, healthy *Traf7^fl/fl*^* mice carried only one allele with *Rik* deletion (lane 8), while both *Traf7^fl*/fl*^* and *Rik^null^* mice displayed a similar runt phenotype and were homozygous for the *Rik^null^* allele (lanes 9-10). Mice heterozygous for *Rik* deletion were phenotypically indistinguishable from their WT littermates, indicating that *Rik^null^* allele was a recessive carrier of the runt phenotype. Homozygous *Rik^null^* mice were born at expected Mendelian inheritance distribution ratio. They displayed no obvious abnormalities at birth and, like the original runt *Traf7^fl*/fl*^* mice, *Rik^null^* pups gained weight similar to WT littermates until P12 (Figure 1C), when usually the eyes are open, fur growth is complete, and teeth are erupted. After this time point, WT pups progressively gained weight and grew in size by eating more solid food (JAX Mice Pup Appearance by Age (nih.gov). *Rik^null^* mice had the normal numbers of teeth and eyes, but started to lose weight, exhibited ataxic gait, and died around P24, one week earlier than runt *Traf7^fl*/fl*^* mice (Figure 1D, 1E, and Movie 2). In contrast, *Traf7^fl/fl^* mice without *Rik* deletion were healthy and fertile as described in Tsitsikov et al. (Tsitsikov et al., 2023).

### P14 *Rik^null^* brains have an abnormal transcriptional profile

Because the *Rik^null^* mouse phenotype was consistent with CNS abnormalities, we compared brain transcriptome profiles from 14 days old littermate WT and *Rik^null^* pups to gain insight into the developmental differences at the earliest time point of the weight gain bifurcation. The PCA plot of the RNA-seq analysis displayed that both control and *Rik^null^* samples did not form compact clusters (Figure S3A). However, there was a good separation between them, with only one of the eight *Rik^null^* samples located near WT controls. The vast majority of identified genes (13,623) were common between the groups, with only 317 and 230 DEGs upregulated in WT or *Rik^null^* brains, respectively (Figure S3B). The Heat map revealed high sample similarity within each group and a good separation between the groups (Figure S3C).

Remarkably, the expression of *Rik* in *Rik^null^* brains decreased more than 8-fold and showed the greatest difference in relative expression compared to WT counterparts (Figure 2A). In contrast, the expression of closely positioned neighboring genes *alpha-tubulin N-acetyltransferase 1* (*Atat1*) and *pre-mRNA-splicing factor ATP-dependent RNA helicase* (*Dhx16*) was not inhibited (Supplemental file 2), indicating that the 59-bp deletion only interfered with *Rik* expression and, therefore, might contain gene-specific regulatory elements important for its transcription.

**Figure 2.**
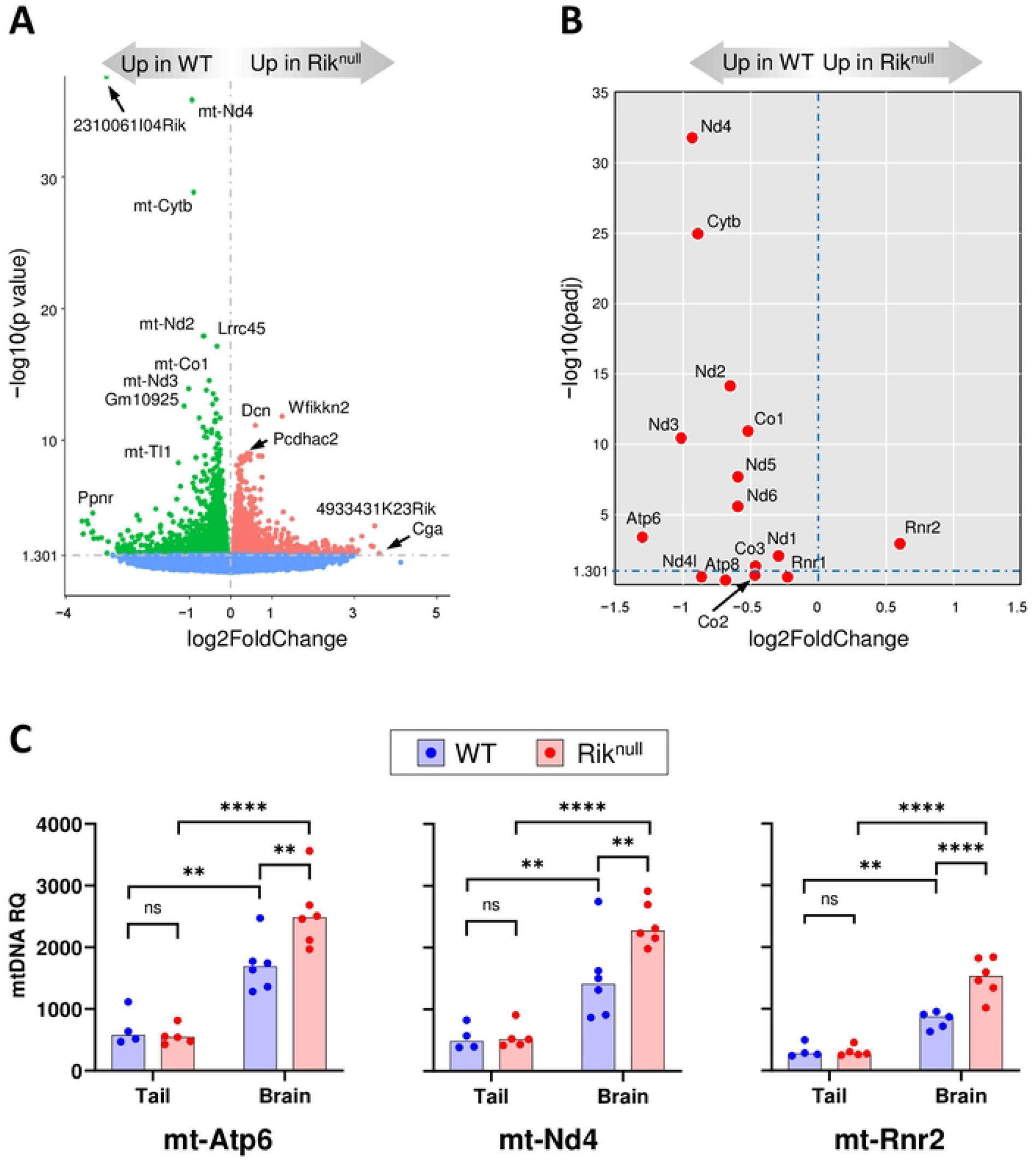
Analysis of 14 days old *Rik^null^* brain transcriptome. (**A**) Volcano plot of RNA-seq analysis visualizing significant DEGs in P14 *Rik^null^* versus WT brains. Magnitude of change (x-axis) vs. statistically significant *p* values (y-axis). Points that have a *p* value less than 0.05 (−log10=1.301) are shown in blue (n=6 WT, n=5 *Rik^null^*). (**B**) Scatter plot of RNA expression levels from 13 messenger and 2 ribosomal RNA genes located in mitochondrial DNA: *Rik^null^* versus WT relative expression (x-axis) vs. statistical significance (y-axis) of the difference. (**C**) Relative quantity (RQ) of mtDNA in brains and tails in WT and *Rik^null^* mice. RQ of three representative mitochondrial genes were measured by gene-specific quantitative real time PCR using nuclear DNA-coded *Gapdh* gene as internal control. Two-factor (genotype, tissue) ANOVA with Bonferroni post-hoc test. n=4-6 WT, n=5-6 *Rik^null^*. ** p < 0.01, *** p < 0.001, **** p < 0.0001, ns - non-significant.

Five of the top seven DEGs with the highest significant difference between *Rik^null^* and control mice were genes encoded by mitochondrial DNA, including *mt-Nd4*, *mt-Cytb*, *mt-Nd2*, *mt-Co1*, and *mt-Nd3* (Figure 2A). Because these genes encode subunits of three mitochondrial proton pumps, we examined the expression of all 13 messenger and two ribosomal RNAs (Figure 2B). Of the 15 genes, 14 showed reduced expression in *Rik^null^* brains, with only *mt-Rnr2*, which encodes a large subunit of ribosomal 16S RNA, showing increased expression. Thus, the *Rik^null^* mutation caused a substantial reduction in the expression of mitochondrial genes. While the highest statistical significance was observed for *mt-Nd4*, a subunit of mitochondrial respiratory chain complex I, the biggest decline in expression was displayed by *mt-Atp6*, encoding a subunit of the F_1_F_0_ATP-synthase complex (Figure 2B). Accordingly, gene set enrichment analysis in the Kyoto Encyclopedia of Genes and Genomes (KEGG) pathway database unveiled higher expression of thermogenesis and reactive oxygen species genes in WT controls compared to mutants (Figure S4A). In contrast, genes associated with the phosphatidylinositol/protein kinase B (PI3-Akt) signaling pathway, human papilloma virus infection, and axon guidance were significantly enriched in *Rik^null^* brains (Figure S4B). The PI3-Akt pathway and axon guidance proteins are important for proper CNS axon growth (Berry et al., 2016). Enriched expression of these genes in the brains of *Rik^null^* mice into the third week of postnatal development, when axonal growth should be mostly completed in altricial rodents such as mice and rats (Zeiss, 2021), is consistent with white matter anomalies noted in *Rik^null^* mice (see below).

To examine whether the difference in brain expression of mitochondrial genes between *Rik^null^* and WT mice is due to different number of mitochondrial DNA (mtDNA) copy number per cell (CN), we looked at mtDNA CN by measuring the relative quantity (RQ) of three representative mitochondrial genes by gene-specific quantitative real time PCR (qPCR). As internal control we used *Gapdh*, which is coded by nuclear DNA and presented in two copies in each mononucleated cell. We have chosen *mt-Atp6*, *mt-Nd4*, and *mt-Rrn2*, because the first two genes demonstrated decreased mRNA expression in *Rik^null^* brains compared to WT counterparts, while the third one displayed the opposite result, an increased expression in *Rik^null^* brains compared to controls (Figure 2B). We also compared the RQ of mtDNA between tails and brains in the same groups of mice. The experiments revealed that mtDNA of each of these genes was no different between *Rik^null^* and WT tails (Figure 2C). In contrast, *Rik^null^* brains demonstrated significantly higher mtDNA content compared to WT brains and this observation was true for all three genes, independently of mRNA expression levels of each of these genes. Thus, these results suggested that P14 *Rik^null^* brains have higher mtDNA copy numbers compared to controls. Interestingly, there were significantly higher RQ of mtDNA in the brains compared to tails in *Rik^null^* mice as well as in WT animals.

### Low expression of genes specific for myelinating oligodendrocytes in P23 *Rik^null^* brains

Next, we compared transcriptomes of WT and *Rik^null^* brains at P23, the peak of the difference between the groups. RNA-seq analysis of P23 *Rik^null^* brains revealed that the groups were well separated on PCA plot (Figure S5A). There were 452 and 601 DEGs expressed in WT or *Rik^null^* brains, respectively (Figure S5B). Heat map analysis demonstrated high sample similarity within each group and a good separation between the groups (Figure S5C). As expected, *Rik* showed the greatest difference in relative mRNA expression between WT and *Rik^null^* brains (Figure 3A). As on P14, all mitochondrial messenger RNAs exhibited significant enrichment in WT brains compared to *Rik^null^* brains (Figure 3B). Like the results above, *mt-Rnr2* displayed increased expression in *Rik^null^* brains compared to WT controls, while the expression of *mt-Rnr1* did not differ between groups. It is interesting that *transferrin* (*Trf*), which encodes a critical iron transporter, displayed one of the highest differences in expression between *Rik^null^* WT control brains (Figure 3A). *Rik^null^* brains also expressed lower levels of genes of two enzymes responsible for biosynthesis of creatine, *glycine amidinotransferas*e (*Gatm*) and *guanidinoacetate N-methyltransferase* (*Gamt*). Of note, oligodendrocytes (OLs) exhibit the highest average level of both *Gatm* and *Gamt* mRNA expression across the entire body (Baker et al., 2021). KEGG pathway enrichment analysis revealed reduced expression of several genes associated with neurodegenerative pathways in *Rik^null^* brains (Figure S6A). In contrast, there was elevated expression of genes associated with the cytokine-cytokine receptor interaction pathway in *Rik^null^* brains (Figure S6B).

**Figure 3.**
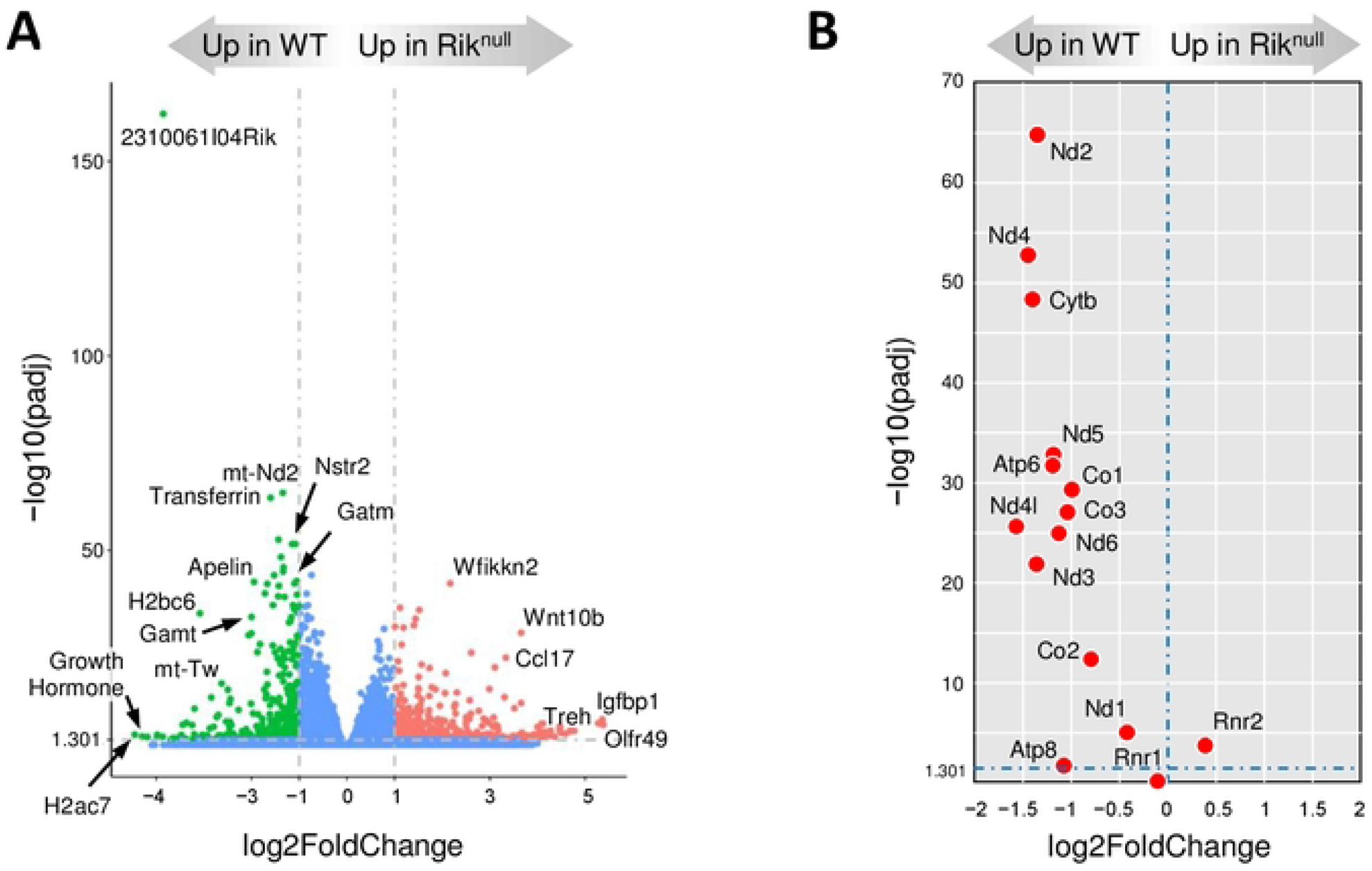
Analysis of 23 days old *Rik^null^* brain transcriptome. (**A**) Volcano plot of RNA-seq analysis visualizing significant DEGs in P23 *Rik^null^* versus WT (control) brains. Magnitude of change (x-axis) vs. adjusted *p* values (padj) (y-axis). Points that have a fold change less than 2 (log_2_=1) or have a *padj* value less than 0.05 (-log10=1.301) are shown in blue (n=6 WT, n=8 *Rik^null^*). (**B**) Scatter plot of mitochondrial gene expression levels for 13 protein-coding and 2 ribosomal RNA genes: *Rik^null^* versus WT relative expression (x-axis) vs. statistical significance (y-axis) of the difference.

To ascertain whether the differences in transcriptome profiles between *Rik^null^* and WT brains could be attributed to specific cell populations, we compared the expression of the most highly expressed genes (excluding transcription factors) specific to several key cell populations in the brain (Zhang et al., 2014). The analysis revealed no significant differences in the expression of the selected neuronal, endothelial, and pericyte marker genes (Figure 4A and Supplemental files 3 and 4). The only microglia-specific DEG showing significantly lower expression in *Rik^null^* brains compared to WT brain was *Selectin P Ligand* (*Selplg*, also called *CD162*), suggesting that microglia were relatively unaffected in the *RIK^null^* brain. Four out of six astrocyte-specific genes displayed lower expression in *Rik^null^* brains compared to WT brains. These genes included *aquaporin 4* (*Aqp4*) and *metabotropic glutamate receptor 3* (*Grm3*), suggesting dissimilar astrocyte transcriptome profiles in *Rik^null^* and WT brains. Interestingly, OL precursor cells (OPCs) had two significant DEGs, *Mmp15* and *Rlbp*, while the expression of two other well-known OPC marker genes, *platelet-derived growth factor receptor α (Pdgfrα*) and *chondroitin sulfate proteoglycan 4* (*Cspg2,* also called *Ng2*) (Fletcher et al., 2021), did not differ between *Rik^null^* and WT brains (Figure 4A and Supplemental file 4). Gene expression by newly formed OLs (NFOs) did not differ between *Rik^null^* and WT brains. In contrast, six genes specific to myelinating OLs (MOs) showed lower expression in *Rik^null^* brains compared to WT brains, suggesting potential dysfunction in MOs. Importantly, these DEGs included *Opalin*, a gene encoding a CNS myelin marker specifically expressed in MOs that has been shown to be important for fine-tuning of exploratory behavior (Yoshikawa et al., 2016).

**Figure 4.**
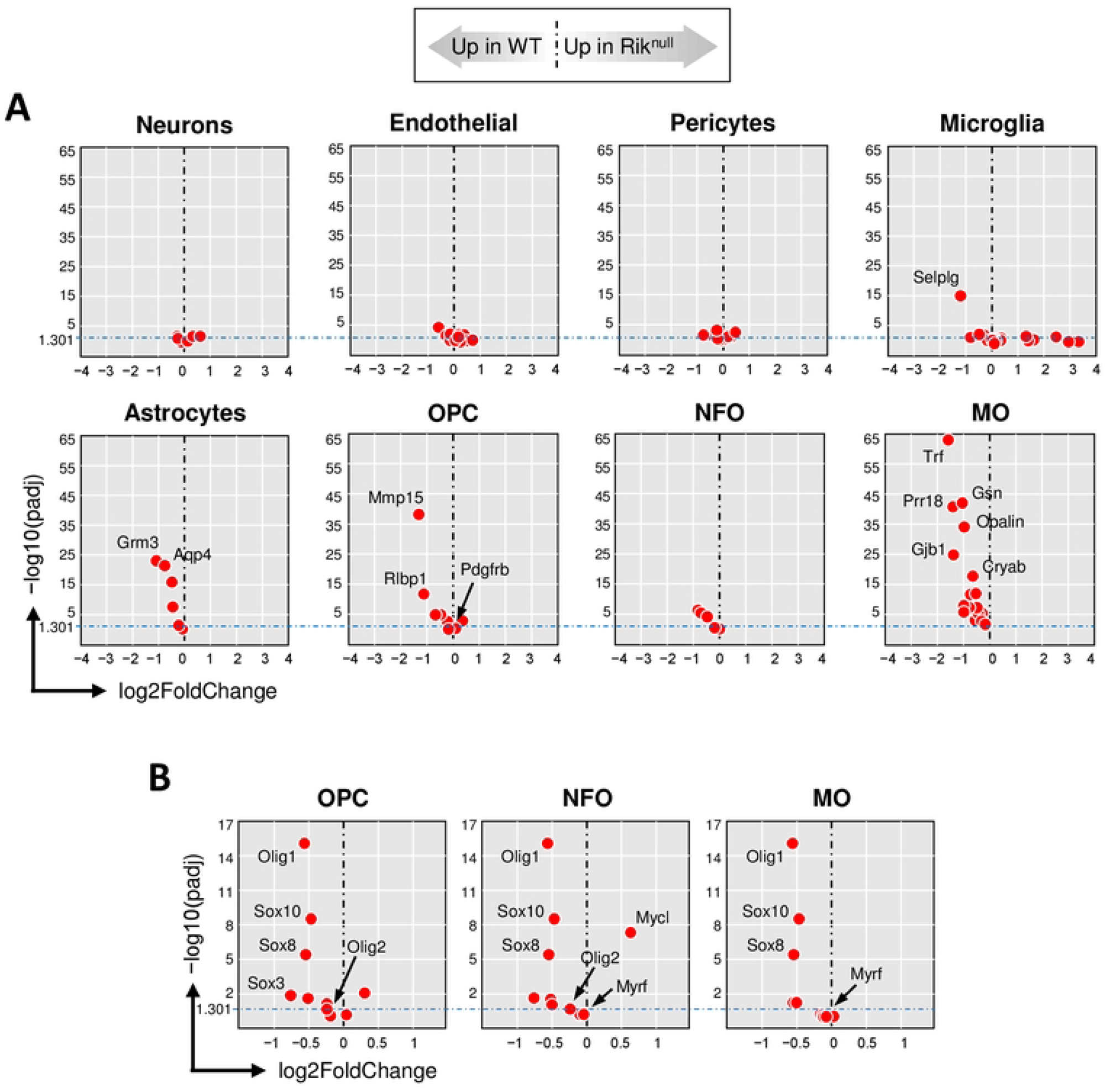
Low expression of genes specific for myelinating oligodendrocytes in 23 days old *Rik^null^* mouse brains. Scatter plots of mRNA expression levels of gene markers specific for different cell populations in in P23 *Rik^null^* versus WT mouse brains from RNA-seq DE analysis: relative expression (x-axis) vs. statistical significance (y-axis) of difference in mRNA expression. OPC: oligodendrocyte precursor cells; NFO: newly formed oligodendrocytes; MO: myelinating oligodendrocytes. (**A**) Cell lineage-specific marker genes. (**B**) Marker genes coding transcription factors in oligodendrocyte populations.

Next, we compared the expression of oligodendrocyte-specific transcription factors in *Rik^null^* and WT brains (Zhang et al., 2014). Pan-OL transcription factors *oligodendrocyte transcription factor 1* (*Olig1*), and *SRY-box transcription factors* (*Sox8* and *Sox10*) differed significantly and showed lower expression in *Rik^null^* OLs compared to WT OLs (Figure 4B and Supplemental file 5). In contrast, the expression of *Olig2* (expressed only in OPC and NFO populations) did not differ significantly between *Rik^null^* and WT brains. Moreover, *myelin regulatory factor* (*Myrf*) (expressed only in NFO and MO populations) was expressed to the same extent in *Rik^null^* and WT brains. Taken together, these results suggested that *Rik* is potentially important for differentiation of NFOs into MOs in the mouse brain during the third week of postnatal development.

### Discrete myelination of anterior thalamic nuclei in P23 *Rik^null^* brains

Under visual examination, the brain of P23 mutant mice appeared without gross deformities, except that the whole brain, and particularly the cortical hemispheres, appeared consistently smaller compared to WT controls (Figure S7A). To understand the consequences of *Rik* deficiency for CNS myelination, we used the myelin stain Luxol Fast Blue (LFB) to stain *Rik^null^* brains. Of note, LFB stains myelin fibers in vibrant blue, Nissl substance in dark blue, and cell nuclei in blue. The LFB staining intensity in *Rik^null^* brain sections was reduced compared to WT sections (top panels in Figure 5A), suggesting decreased total myelination of *Rik^null^* brains compared to their WT counterparts. Anterior thalamic nucleus (ATN) was one of the most affected regions (bottom panels in Figure 5A). While the staining intensity of anterodorsal nuclei (ADN) in *Rik^null^* and WT brains was similar, the intensity of *Rik^null^* anteroventral nuclei (AVN) appeared much weaker compared WT controls. At higher magnification, *Rik^null^* and WT and demonstrated similar intensity of longitudinal axon fibers (top panels in Figure 5B). In contrast, cross-sectioned axons in *Rik^null^* AVN displayed decreased LFB-staining intensity compared to WT counterparts (bottom panels in Figure 5B). Moreover, the cytoplasm of *Rik^null^* cells appeared much smaller and contained less Nissl substance, indicating decreased protein biosynthesis compared to WT OLs. Haematoxylin and Eosin staining (H&E) of those regions demonstrated no obvious variations in cellularity or general structure between mutant and control ATNs (Figure S7B).

**Figure 5.**
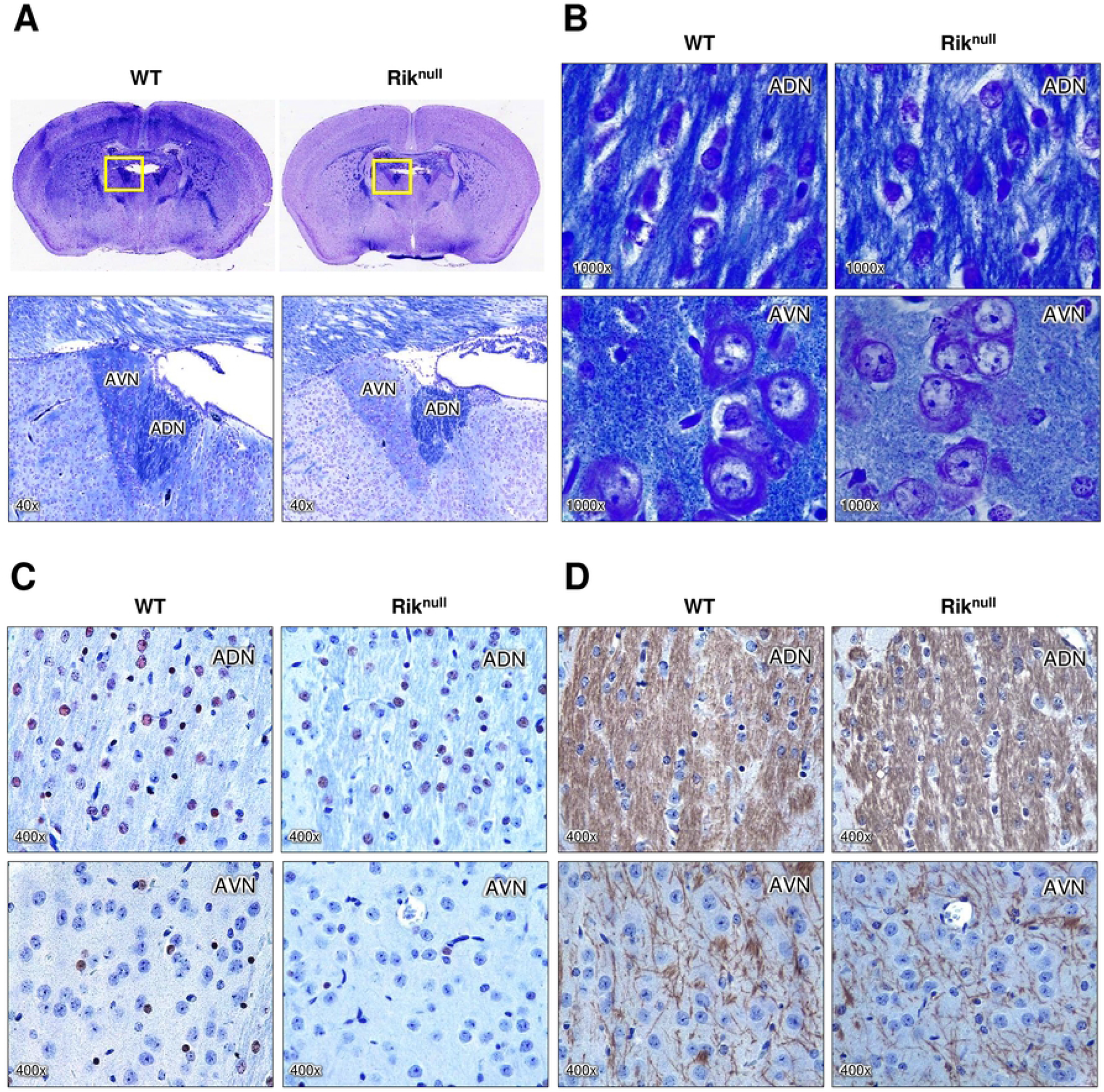
RIK deficiency causes defects in myelinated tracts of the anterior thalamic nuclei. (**A**) Luxol Fast Blue (LFB) staining of brain coronal sections from 23 days old WT and *Rik^null^* mice. Top panels are representative of brain sections at Bregma −0.58 mm. Bottom panels show magnified images of the area within yellow windows above containing anteroventral nuclei (AVN) and anterodorsal nuclei (ADN). (**B**) Representative views of LFB-stained ADN and AVN at higher magnification as marked. (**C** and **D**) Hematoxylin and immunohistochemistry staining of ADN and AVN regions with oligodendrocyte-specific anti-Olig2 (C) and anti-MBP (D) antibodies. Brown color indicates corresponding protein location. Cell nuclei are colored blue.

There were no clear differences in distribution and density of OLs in ADN between *Rik^null^* and WT brains, as determined by immunolabeling for the pan-OL marker Olig2 (top images in Figure 5C). Interestingly, the number of OLs in AVNs seems lower than the number of OLs in ADNs in WT mice as well as *Rik^null^* animals (compare top to corresponding bottom images in Figure 5C). The number of OLs in AVN of *Rik^null^* mice was even lower than WT AVNs. In agreement with LFB staining of ADNs, staining with anti-MBP antibodies revealed similar myelination of these nuclei in *Rik^null^* and WT mice (top panels in Figure 5D). The density of myelinated fibers appeared remarkably lower in AVNs compared to ADNs in both *Rik^null^* and WT mice (compare top to corresponding bottom images in Figure 5D). Furthermore, MBP staining of *Rik^null^* AVNs was less intense than in WT controls, suggesting that RIK is important for myelination of axons in AVNs during late postnatal development.

### Decreased myelination of anterior commissure (AC) in P23 *Rik^null^* brains

Further analysis revealed that the most dramatic divergence between control and mutant brains occurred in AC (Figure 6). *Rik^null^* AC displayed much weaker LFB staining compared to WT AC. The difference was apparent in both coronal and as well as sagittal views (Figure 6A and C). Similar to other mouse models with myelination defects (Menichella et al., 2003, Neusch et al., 2001), numerous vacuoles of varying size were detected in AC of *Rik^null^* mice, but not in WT mice (Figure 6A and C). The extensive vacuolation was also evident with H&E staining (Figure 6B and D).

**Figure 6.**
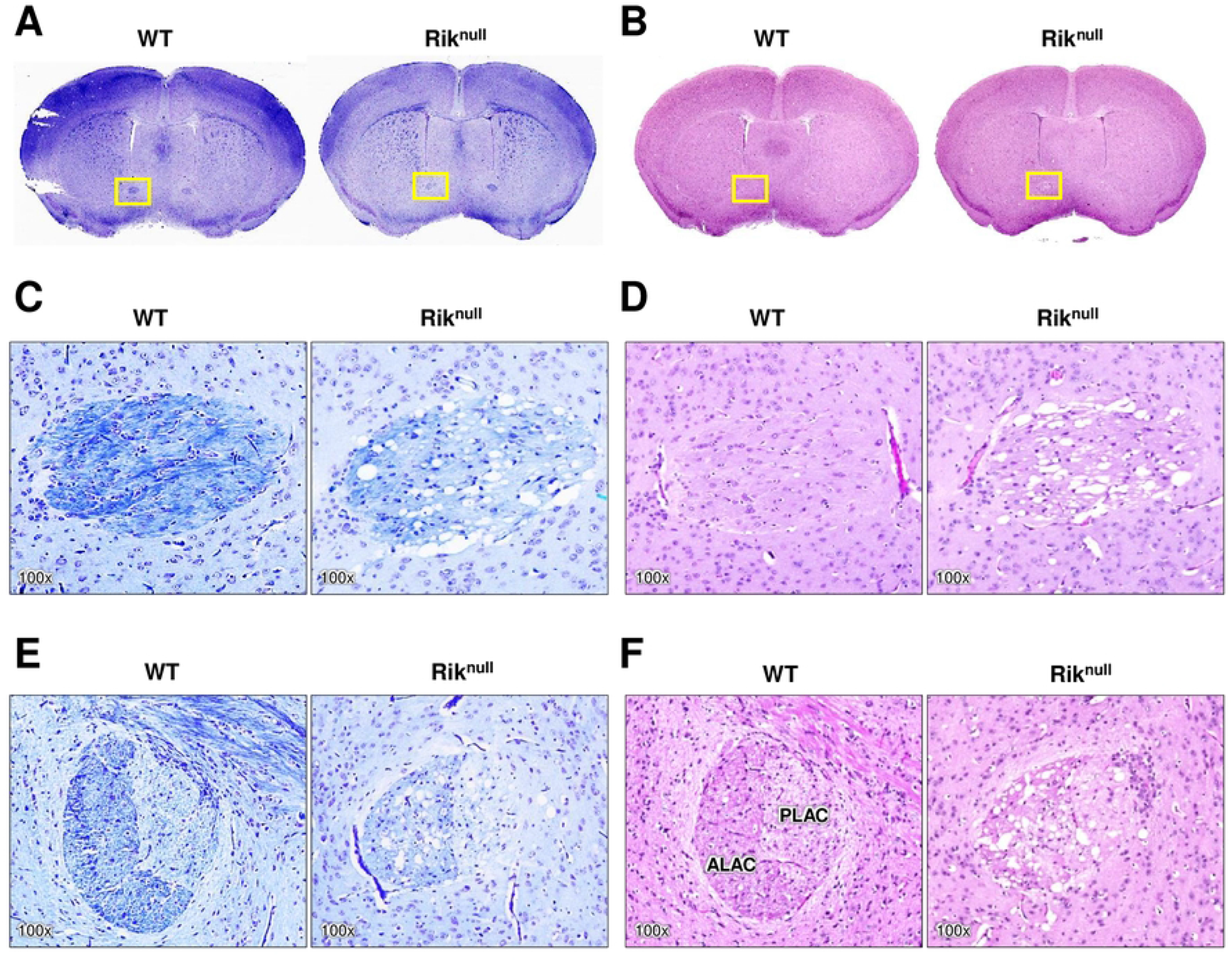
RIK deficiency causes defects in myelinated tracts of anterior commissure. Histological investigation of brain coronal sections from 23 days old mice at Bregma 0.38 mm. WT and *Rik^null^* brain sections were stained with LFB (A and C) or H&E (B and D). (**C** and **D**) Insets of yellow windows in panels (**A** and **B**) above at higher magnification. (**E** and **F**) Magnified image of anterior commissure on sagittal sections (not shown here) from P23 mice (∼lateral 0.24 mm). ALAC: anterior limb of the anterior commissure; PLAC: posterior limb of the anterior commissure. Total magnification of images is marked on each image.

To assess the potential cellular basis for the histological observations, we stained for cell lineage specific markers. Staining with antibodies to glial fibrillary acidic protein (GFAP), a major marker of mature astrocytes in CNS, showed the expected complex astrocytic morphology in the *Rik^null^* brain (Figure 7A and 7B), suggesting no obvious abnormalities in astrocyte development, in agreement with our RNA-seq results (Figure 4A). However, the ramification of fine processes of *Rik^null^* astrocytes looked more disorganized compared to WT counterparts. Staining for OLIG2 revealed comparable numbers of OLs in *Rik^null^* and control ACs, while distribution of *Rik^null^* OLs looked more chaotic compared to WT counterparts, which mostly aligned along the axonal fibers (Figure 7C and 7D). Immunolabeling with antibodies against proteolipid protein 1 (PLP1) and myelin basic protein (MBP) showed a reduction of myelin sheaths in the AC of *Rik^null^* mice compared to controls (Figure 7E, F, G and H). Mature OLs and their myelin sheaths, identified by Opalin immunolabeling (Figure 7I and 7J), showed more intense labeling in the WT AC than in the AC of *Rik^null^* mice. Interestingly, the intensity of staining of OL cytoplasm in AC of *Rik^null^* mice was stronger than the myelin sheaths.

**Figure 7.**
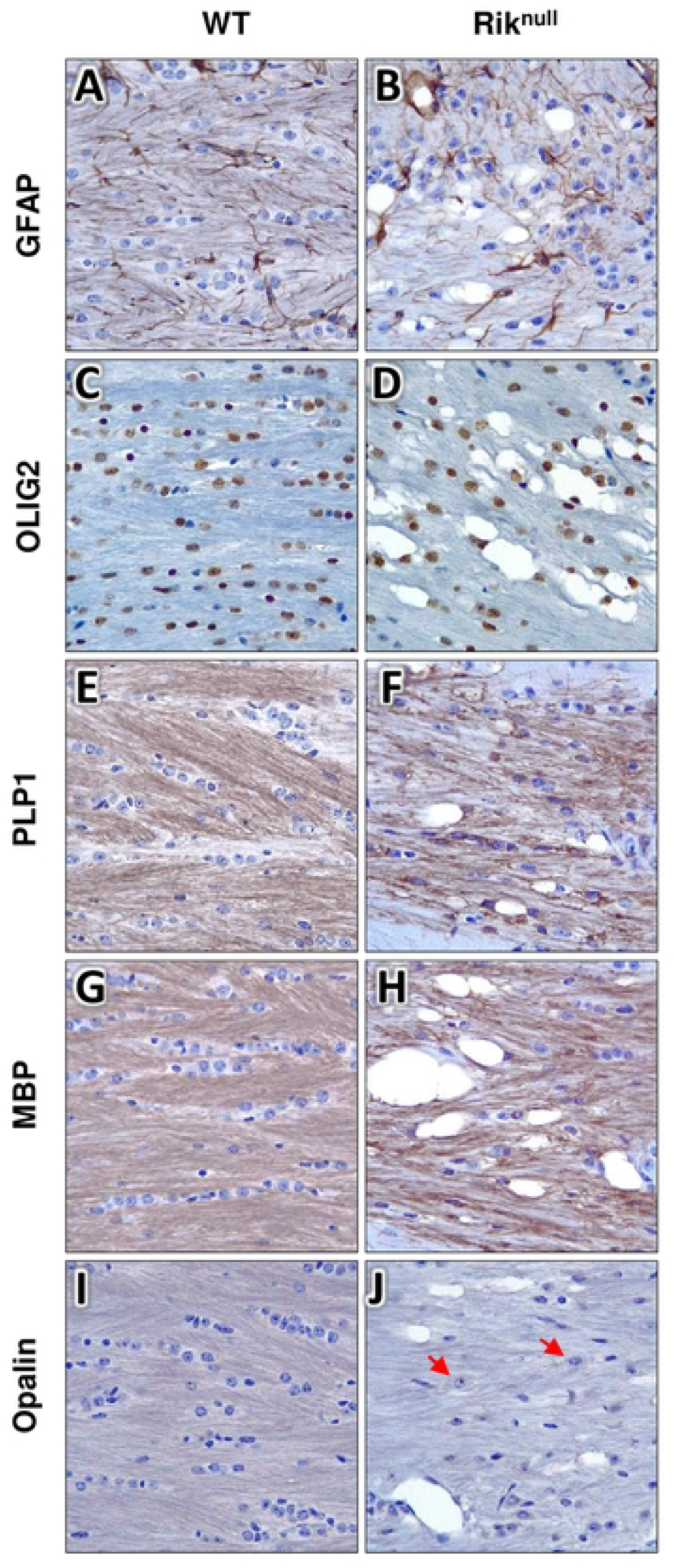
Immunohistochemical (IHC) analysis of 23 day old WT and *Rik^null^* mouse brains. Representative IHC with hematoxylin staining of WT and *Rik^null^* mouse brains at P23 with anti-GFAP antibody, an astrocyte-specific marker (A and B) or antibodies against oligodendrocyte-specific markers as labelled on images (panels C-J). Total magnification of all shown images is 400x. Regions of coronal sections at Bregma −0.58 mm show parts of anterior commissure heavily myelinated in WT as shown in Figure 6C.

## Discussion

In this study, we demonstrate that a 59-bp homozygous deletion in the first intron of the *2310061I04Rik* gene abolishes its expression. Mouse *Rik* is predicted to encode five protein isoforms of different length with a 315 amino acid long canonical isoform (Uniprot.org entry #B8JJ69). RIK protein is evolutionarily conserved from *C. elegans* to mammals (https://useast.ensembl.org/Homo_sapiens/Gene/Compara_Tree?db=core;g=ENSG00000204564;r=6:30647039-30653210). The human orthologue of RIK is called C6orf136. Mammalian RIK proteins consist of two domains: a proline-rich domain and a domain of unknown function 2358 (duf2358). RIK has no recognized function. However, it is probably unique since there are no RIK paralogues in the human or mouse genome. RIK was found to localize to mitochondria (https://personal.broadinstitute.org/scalvo/MitoCarta3.0/mouse.mitocarta3.0.html), indicating that RIK might be important for mitochondria activity.

When the *Rik^null^* phenotype was first observed on the *Traf7^fl/fl^* background (Figure S1), the weight of runt *Traf7^fl*/fl*^* mice reached a maximum of 7g, and they died before P40. *Rik^null^* mice carrying wildtype *Traf7* after backcrossing into C57BL6/J strain reached the same weight, but the majority of them died before P25, substantially earlier than runt *Traf7^fl*/fl*^* mice. These discrepancies in life expectancy between mutant *Rik^null^* and *Traf7^fl*/fl*^* mice with *Rik* deletion may be due to complex multigenic differences among commonly used C57BL/6 substrains (Yoshiki and Moriwaki, 2006, Mekada and Yoshiki, 2021). They may result from differences among genes located between *Rik* and the centromere of the telocentric chromosome 17 (Figure 1A), which interestingly includes the whole *Qki* locus.

The human mitochondrial genome is a single circular double-stranded DNA molecule of almost 17 kilobase pairs (Jedynak-Slyvka et al., 2021). MtDNA contains 37 genes that encode 13 proteins, two ribosomal RNAs, and 22 transfer RNAs. Our RNA-seq experiments demonstrated low expression of most mitochondrial genes in P14 and P23 *Rik^null^* brains. Of the 15 genes, encoding all messenger and ribosomal RNAs, 14 showed reduced expression, while only *mt-Rnr2* demonstrated increased expression in *Rik^null^* brains compared to WT controls (Figure 2 and 3). On the other hand, mtDNA copy numbers were increased in *Rik^null^* brains compared to controls (Figure 2C), indicating that *mt-Rrn2* was the only gene expressed in agreement with the mtDNA copy number in *Rik^null^* brains, while the lower expression of other mitochondrial messenger RNAs and mt-Rrn1 contradicted the higher mtDNA copy numbers in *Rik^null^* brains. Since all 15 mentioned genes, except *mt-ND6* are transcribed from the same heavy strand promoter, these observations suggest that RIK regulates the stability of mitochondrial RNAs. Future studies are needed to investigate molecular mechanisms of RIK function.

*Rik^null^* mouse brain showed lower expression of genes encoding subunits of all electron transport pumps, including respiratory complex I (*mt-Nd2*, *mt-Nd4* and others), complex III (*mt-Cyb*), and complex IV (*mt-Co1*). It is interesting that, similarly to *Rik^null^* animals, mice deficient in a subunit of mitochondrial complex I, *NADH dehydrogenase [ubiquinone] iron-sulfur protein 4* (*Ndufs4*), were born apparently healthy (Kruse et al., 2008) Three weeks later, they lagged behind in growth, developed ataxia and died early, similar to *Rik^null^* mice. However, *Ndufs4* knockout mice reached their maximum body weight of 15g at P28, which is two weeks later than *Rik^null^* animals. At that age, the vast majority of *Rik^null^* mice were already dead. *Rik^null^* mice never survived beyond 4 weeks of age in contrast to *Ndufs4* knockout mice that survived up to P50, which is much longer than survival by *Rik^null^* animals. *Ndufs4* deficient mice were developed to study inherited mitochondrial dysfunction disorders, because mutations in the nuclear DNA-encoded *Ndufs4* gene in humans induce ‘mitochondrial complex I deficiency, nuclear type 1’ (MC1DN1) and Leigh syndrome in children (van de Wal et al., 2022). Since genotype to phenotype translation depends on genetic environment of any given allele, these differences between *Ndufs4* knockout and *Rik^null^* mice may result from different genetic backgrounds. *Ndufs4* knockout mice were on a mixed 129/Sv x C57BL/6 genetic background, while *Rik^null^* mice were on C57BL/6 background.

P23 *Rik^null^* brains demonstrated lower expression of GATM and GAMT, which function together to synthesize creatine to facilitate recycling of ATP by converting ADP back to ATP. Mature OLs are the major producers of creatine during postnatal development (Rosko et al., 2023) and express higher levels of both enzymes compared to any other brain cell type, including OPCs (Baker et al., 2021). Although *Gatm-* or *Gamt-*deficient mice consistently weighed less than control littermates, they survived until 24 weeks of age and beyond (Rosko et al., 2023, Choe et al., 2013, Schmidt et al., 2004). Overall, the phenotype of *Gatm* or *Gamt-* deficient mice was much milder compared to *Rik^null^* mice. Therefore, a decreased expression of *Gatm* and *Gamt* in P23 *Rik^null^* brains suggests that RIK may either directly control their gene expression or indirectly regulate their levels by being important for the process of OL maturation.

P23 *Rik^null^* brains expressed almost 4-fold lower levels of *Trf* compared to WT controls. *Trf*, encodes transferrin,a major transporter of iron, which is an essential nutrient for most forms of life. In addition to the role of transporting oxygen, iron participates in the catalysis of redox reactions throughout the whole body, including the brain. Iron is essential for myelin production as a cofactor for enzymes involved in ATP, cholesterol, and lipid biosynthesis. Furthermore, OLs show the highest iron concentrations in the brain, which is directly linked to their elevated metabolism associated with the high-energy process of myelination (Reinert et al., 2019, Cheli et al., 2020). To be transported through blood and across OL cellular membranes, iron binds to TRF (Abe et al., 2022, Murakami et al., 2019), which is essential for OL maturation and function (Saleh et al., 2003). In fact, OLs are most vulnerable to transferrin deficiency during the premyelinating stage (Espinosa de los Monteros et al., 1999).

It is noteworthy that the genes specific for mature OLs, including *Trf*, *Gatm*, *Gamt,* and *Opalin*, displayed no differences in their expression between P14 *Rik^null^* and WT brains, while the expression of mitochondrial genes was already lower at that time. Considering this, we propose that RIK is important for optimal transcription of mitochondrial genes, specifically in the brain during the third week of postnatal development, when the phenotypic divergence of OL maturation between *Rik^null^* and control mice occurred. During this period of postnatal development, the most rapid phase of myelination begins in the mouse brain (Sturrock, 1980, Nishiyama et al., 2021, Fletcher et al., 2021) and the highest supply of cellular energy is required (Rao et al., 2017). Because our investigation of brains from *Rik^null^* mice revealed that RIK deficiency disturbed the generation of mature OL, we tentatively propose to name RIK as an “oligodendrocyte maturation factor” (OMF).

## Supplemental figure legends

**Figure S1. Severe runting and a shortened life-span of *Traf7^fl/fl^* mice with serendipitous deletion in *2310061I04Rik* gene (*Traf7^fl*/fl*^)*.**

**(A)** Mouse weight chart from P13 to end of life of *Traf7^fl*/fl*^* mice. (n=391 WT, n=480 *Traf7^+/fl*^*, n=230 *Traf7^fl*/fl*^*). Data presented as median±SEM.

**(B)** Representative 4-weeks old WT and *Traf7^fl*/fl*^* littermates.

(**C**) Probability of survival of *Traf7^fl*/fl*^* mice. n=25 *Traf7^fl*/fl*^*

**Figure S2. Differential gene expression in WT and *Traf7^fl*/fl*^* mouse embryos at E9.5.**

**(A)** PCA Plot of RNA-seq analysis in WT and *Traf7^fl*/fl*^* mouse embryos. Each point corresponds to an individual embryo.

**(B)** Venn diagram of RNA-seq analysis in WT and *Traf7^fl*/fl*^* mouse embryos showing DEGs overlap.

**(C)** Heatmap of mRNA expression levels for all significant DEGs in WT and *Traf7^fl*/fl*^* mouse embryos.

**(D)** Volcano plot of RNA-seq analysis visualizing significant DEGs in WT versus *Traf7^fl*/fl*^* embryos: magnitude of change (*x-axis*) vs. statistically significant *p* values (*y*-axis). Points with *p* value less than 0.05 (-log10=1.301) are shown in blue.

**Figure S3. Differential gene expression in 14 days old WT and *Rik^null^* brains.**

**(A)** PCA Plot of RNA-seq analysis in WT and *Rik^null^* brains. Each point corresponds to an individual brain sample.

**(B)** Venn diagram of RNA-seq analysis in *Rik^null^* versus WT brains showing DEGs overlap.

**(C)** Heatmap of mRNA expression levels for all significant DEGs in *Rik^null^* versus WT brains.

**Figure S4. KEGG enrichment analysis of signaling pathways in P14 *Rik^null^* brains.**

Signaling pathways upregulated in WT versus *Rik^null^* brains (A) or in *Rik^null^* versus WT brains (B). Each bubble represents a KEGG pathway. Gene ratio (x-axis) is the proportion of the total genes in a given pathway that is upregulated in the indicated group.

**Figure S5. Differential gene expression in 23 days old WT and *Rik^null^* brains.**

**(A)** PCA Plot of RNA-seq analysis in WT and *Rik^null^* brains. Each point corresponds to an individual brain sample.

**(B)** Venn diagram of RNA-seq analysis in *Rik^null^* versus WT brains showing DEGs overlap.

**(C)** Heatmap of mRNA expression levels for all significant DEGs in *Rik^null^* versus WT brains.

**Figure S6. KEGG enrichment analysis of signaling pathways in 23 days old *Rik^null^* brains.**

Signaling pathways upregulated in WT versus *Rik^null^* brains (A) or in *Rik^null^* versus WT brains (B). Each bubble represents a KEGG pathway. Gene ratio (x-axis) is the proportion of the total genes in a given pathway that is upregulated in the indicated group.

**Figure S7. Morphological comparison of WT and *Rik^null^* brains.**

**(A)** Images of a dorsal view of the brains from 3 week old WT and *Rik^null^* littermates.

**(B)** Coronal sections of brain shown in (A) stained with hematoxylin and eosin. Top panels are representatives of brain sections at about Bregma −0.58 mm. Bottom panels show magnified images of the area within black windows above containing anteroventral nuclei (AVN) and anterodorsal nuclei (ADN).

## Materials and Methods

### Discovery of Rik^Null^ mice

All housing and experimental use of mice were carried out in AAALAC-accredited facility in accordance with United States federal, state, local, and institutional regulations and guidelines governing the use of animals and were approved by OUHSC Institutional Animal Care and Use Committee. The *Rik^Null^* allele arose as a spontaneous recessive mutation in the *2310061I04Rik* gene during generation of conditional *Traf7* knockout mice. The *Traf7* targeting vector and *Traf7^+/fl^* mice were generated by the Ingenious Targeting Laboratory (Ronkonkoma, NY, USA) on the C57BL/6 background. Since *Rik* is located in proximity of *TRAF7^fl^* allele on chr17, we unlinked *TRAF7^fl^* and *Rik^Null^* alleles by crossbreeding of *Traf7^+/fl*^Rik^+/Null^* littermates until we obtained several mice without pathology. The resulting *Rik^Null/+^* mice were backcrossed on the C56BL/6 background for several generations and were genotyped to confirm presence of *Rik^Null^* allele and absence of *TRAF7^fl^* allele. Mice of both sexes were used for all assays. Mice were humanely euthanized by CO2 asphyxiation immediately prior to tissue collection.

### Tissue DNA preparation

For DNA preparation tissue biopsy was incubated overnight with 0.1 mg/mL proteinase K (Fisher Scientific, Hampton, NH, USA) in ATL buffer at 55°C. Proteins and lipids were precipitated with 1/3 of volume of 5M NaCl and debris was removed by centrifugation at 14,000 rpm for 10 minutes. DNA was precipitated with 7/10 of volume of 100% isopropanol and centrifugation for 10 minutes at 14,000 rpm. The pellet was washed in 70% ethanol and dissolved in TE. DNA concentration was measured with Nanodrop (Fisher Scientific).

### Mouse genotyping

Genotyping of tail DNA was performed on purified genomic DNA using conventional and/or real-time PCR. Genotyping of *TRAF7^fl^* allele was described in (Tsitsikov et al., 2023). Conventional PCR with primers mRik-Fwd: AGATGGAGAAGTCGGTGGA and mRik-Rev: TTTCTCCCATGGAGCAGTAAAC yield amplifies a 401 bp DNA fragment from WT and a 342 bp fragment from 2310061I04Rik^Null^ alleles, respectively. Real-time PCR was performed as a DuPlex reaction for both alleles in one tube using PerfeCTa® qPCR FastMix® II, Low ROX™ (VWR, Radnor, PA, USA, Quanta Biosciences™, #95120) on an Applied Biosystems 7500 Fast Real-Time PCR System (Fisher Scientific). TaqMan Assays for WT and Null alleles are as follows:

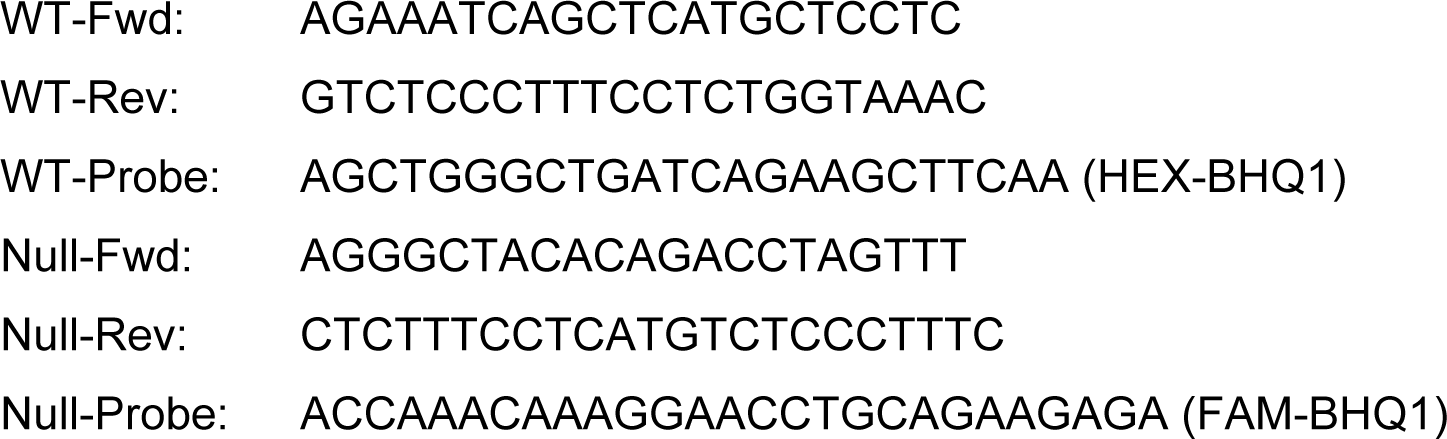

### RNA extraction

Mouse brains were collected and stored in Invitrogen™ RNAlater™ Stabilization Solution (Fisher Scientific, AM7023) before RNA extraction. Total RNA was purified using the RNeasy Lipid Tissue Mini Kit (QIAGEN, Redwood City, CA, USA, 74804) according to the manufacturer’s instructions, quantified by NanoDrop and stored at −80°C.

### RNA-seq and differential expression (DE) analysis

Total brain RNA from at least six WT and 2310061I04RikNull mouse littermates of either sex was used for RNA-seq experiments. Preparation of cDNA libraries, sequencing, and Standard bioinformatics analyses were conducted by Novogene Co., LTD (Beijing, China). Significant DEGs were defined as those that had both an absolute log2FoldChange ≥ 1 as well as a false discovery rate adjusted p-value ≤ 0.05 for each comparison independently.

### Histology

Brains from mice (littermates, different ages, either sex) were fixed in 4% paraformaldehyde, paraffin embedded, and sectioned at 4 μM using standard methods. Sections were stained with either hematoxylin and eosin using Abcam H&E Staining Kit (Fisher Scientific, NC1881153) or Luxol Fast Blue (LFB) using the StatLab LUXOL FAST BLUE KIT (Fisher Scientific, NC9030259) for myelin. Staining was performed according to the manufacturers’ instructions. Images were obtained at total magnifications of 40x and 100x (combination of magnifications of 4x and 10x objective lens with 10x ocular lens) using a Leica (Wetzlar, Germany) DM750 microscope with an ICC50 W Camera Module and included software. Up to seven sections from mice of each genotype were evaluated for each genotype.

### Immunohistochemistry

Slides were deparaffinized in three changes of xylene, two changes of 100 % alcohol, two changes of 95% alcohol, and rehydrated in two changes of H_2_O. Next, slides were bleached with 3% Hydrogen Peroxide (Sigma-Aldrich, St. Louis, Mo, USA, #516813) and masked epitopes were recovered with eBioscience™ IHC Antigen Retrieval Solution - High pH (10X) (Invitrogen, Waltham, MA, USA, #00-4956-58). Slides were blocked for 1 hour at room temperature with blocking solution (1xPBS, 3% milk, 5% Normal Goat Serum, 0.1% Triton X-100, 0.01% NaN3) and incubated with antibodies overnight at 4°C. The antibody dilutions were as follows: GFAP (D1F4Q) XP Rabbit mAb (Cell signaling, Danvers, MA, USA, #12389) (1:500); PLP1 (E9V1N) Rabbit mAb (Cell signaling #28702) (1:1000); MBP (D8X3Q) XP Rabbit mAb (Cell signaling #78896) (1:2000); Opalin polyclonal rabbit antibody (Sigma-Aldrich #HPA014372) (1:1000) and Olig2 polyclonal rabbit antibody (Sigma-Aldrich, #AB9610) (1:200). Slides were washed in three changes of 1xPBS and incubated in SignalStain® Boost Detection Reagent (HRP, Rabbit, Cell signaling #8114) for 1 hour at room temperature. Bound antibodies were revealed with Metal Enhanced DAB Substrate Kit (Thermo fisher, Waltham, MA, USA, #34065). Slides were counterstained with Hematoxylin for 2 minutes and Bluing Reagent for 15 seconds.

### Mitochondrial copy number (mtDNA-CN) measurement

The mtDNA-CN was determined using a duplexed real time qPCR ΔΔC_T_ assay. The cycle threshold (C_T_) value of three mitochondrial genes (*Atp6*, *Rnr2*, and *Nd4*) and nuclear-specific (*Gapdh*) target genes were determined in several replicates for each sample. Relative measure of mtDNA-CN is reported as the difference in C_T_ values (ΔC_T_) for each pair of genes. The mtDNA-CN is presented as relative quantities (RQ) of mitochondrial DNA and calculated by comparing ΔC_T_ of samples from *Rik^null^* and WT mouse tissues. Assays were run as described in Mouse genotyping section above. TaqMan Assays for *Gapdh* and three mitochondrial genes are as follows:

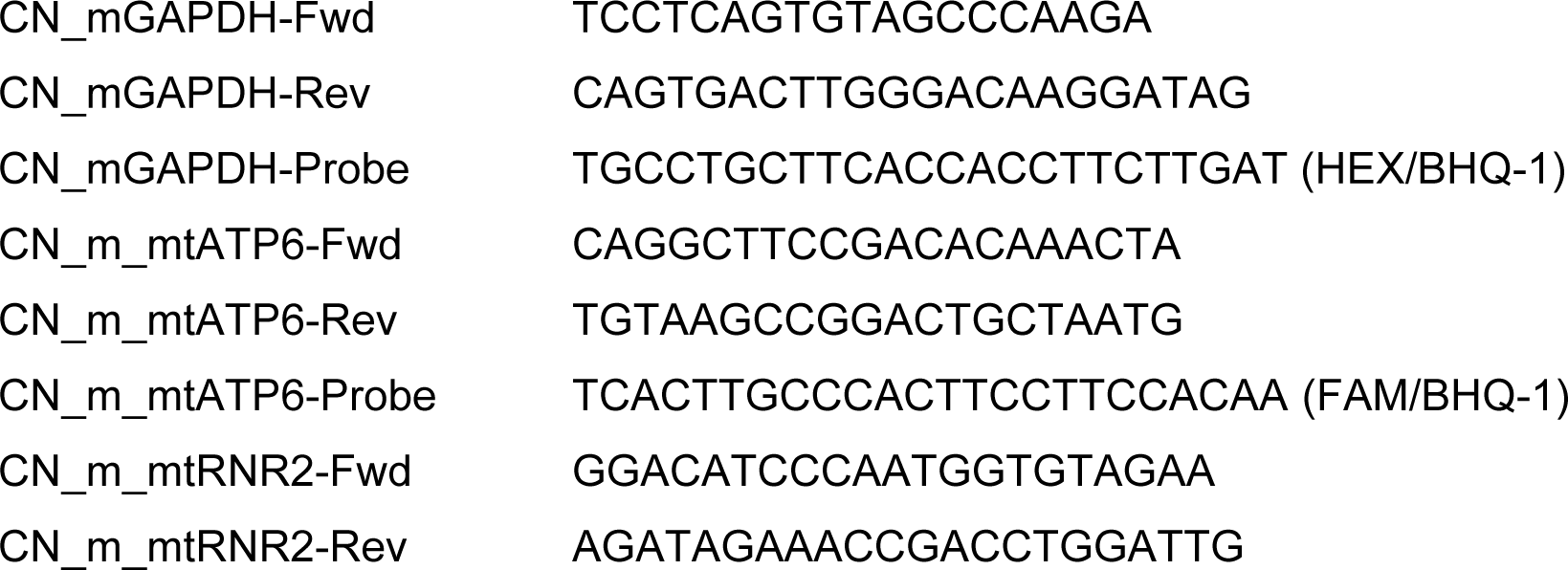
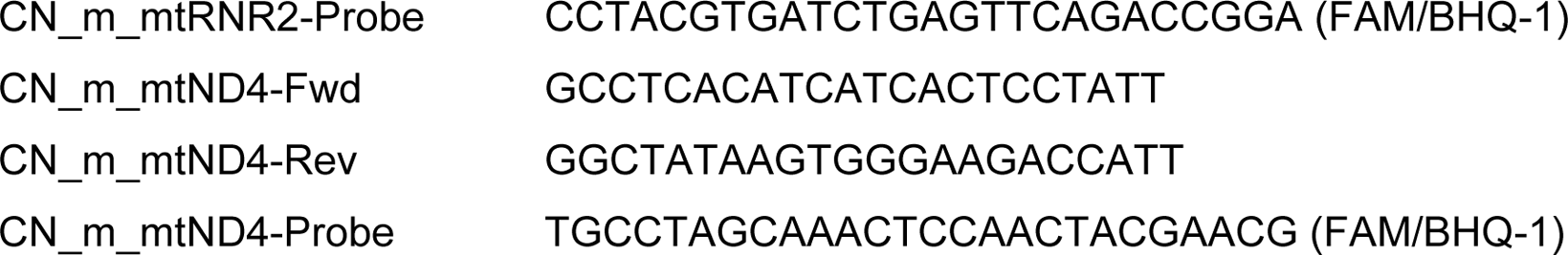

### Statistics

Samples sizes were based on previous studies and limited by availability of genotypes in each litter. Graphpad Prism 9.0 was used for analysis. Data was presented as median± SEM. P < 0.05 was considered statistically significant. Two-factor analysis of variance (ANOVA) with Bonferroni post-hoc analysis was used for Figure 2C. In RNA-seq DE analysis, differential expression was calculated using the Wald test implemented in the R package DESeq2. *mtDNA-CN* were calculated by comparative ΔC_T_ experiment runs on AB7500 Fast machine and analyzed using the 7500 Software v2.3. Calculations of *mtDNA-CN* were performed by Excel with the built-in analysis methods.

## References

Abe, E., Fuwa, T. J., Hoshi, K., Saito, T., Murakami, T., Miyajima, M., Ogawa, N., Akatsu, H., Hashizume, Y., Hashimoto, Y. & Honda, T. 2022. Expression of Transferrin Protein and Messenger RNA in Neural Cells from Mouse and Human Brain Tissue. Metabolites, 12.

Baker, S. A., Gajera, C. R., Wawro, A. M., Corces, M. R. & Montine, T. J. 2021. GATM and GAMT synthesize creatine locally throughout the mammalian body and within oligodendrocytes of the brain. Brain Res, 1770, 147627.

Berry, M., Ahmed, Z., Morgan-Warren, P., Fulton, D. & Logan, A. 2016. Prospects for mTOR-mediated functional repair after central nervous system trauma. Neurobiol Dis, 85, 99–110.

Cheli, V. T., Correale, J., Paez, P. M. & Pasquini, J. M. 2020. Iron Metabolism in Oligodendrocytes and Astrocytes, Implications for Myelination and Remyelination. ASN Neuro, 12, 1759091420962681.

Choe, C. U., Nabuurs, C., Stockebrand, M. C., Neu, A., Nunes, P., Morellini, F., Sauter, K., Schillemeit, S., Hermans-Borgmeyer, I., Marescau, B., Heerschap, A. & Isbrandt, D. 2013. L-arginine:glycine amidinotransferase deficiency protects from metabolic syndrome. Hum Mol Genet, 22, 110–23.

Ebersole, T. A., Chen, Q., Justice, M. J. & Artzt, K. 1996. The quaking gene product necessary in embryogenesis and myelination combines features of Rna binding and signal transduction proteins. Nat Genet, 12, 260–5.

Espinosa De Los Monteros, A., Kumar, S., Zhao, P., Huang, C. J., Nazarian, R., Pan, T., Scully, S., Chang, R. & De Vellis, J. 1999. Transferrin is an essential factor for myelination. Neurochem Res, 24, 235–48.

Fletcher, J. L., Makowiecki, K., Cullen, C. L. & Young, K. M. 2021. Oligodendrogenesis and myelination regulate cortical development, plasticity and circuit function. Semin Cell Dev Biol, 118, 14–23.

Hardy, R. J., Loushin, C. L., Friedrich, V. L., Jr., Chen, Q., Ebersole, T. A., Lazzarini, R. A. & Artzt, K. 1996. Neural cell type-specific expression of Qki proteins is altered in quakingviable mutant mice. J Neurosci, 16, 7941–9.

Haroutunian, V., Katsel, P., Dracheva, S. & Davis, K. L. 2006. The human homolog of the Qki gene affected in the severe dysmyelination “quaking” mouse phenotype: downregulated in multiple brain regions in schizophrenia. Am J Psychiatry, 163, 1834–7.

Hartfuss, E., Förster, E., Bock, H. H., Hack, M. A., Leprince, P., Luque, J. M., Herz, J., Frotscher, M. & Götz, M. 2003. Reelin signaling directly affects radial glia morphology and biochemical maturation. Development, 130, 4597–609.

Hayashizaki, Y. 2003. The Riken mouse genome encyclopedia project. C R Biol, 326, 923–9.

Jedynak-Slyvka, M., Jabczynska, A. & Szczesny, R. J. 2021. Human Mitochondrial RNA Processing and Modifications: Overview. Int J Mol Sci, 22.

Kruse, S. E., Watt, W. C., Marcinek, D. J., Kapur, R. P., Schenkman, K. A. & Palmiter, R. D. 2008. Mice with mitochondrial complex I deficiency develop a fatal encephalomyopathy. Cell Metab, 7, 312–20.

Lakso, M., Pichel, J. G., Gorman, J. R., Sauer, B., Okamoto, Y., Lee, E., Alt, F. W. & Westphal, H. 1996. Efficient in vivo manipulation of mouse genomic sequences at the zygote stage. Proc Natl Acad Sci U S A, 93, 5860–5.

Mekada, K. & Yoshiki, A. 2021. Substrains matter in phenotyping of C57BL/6 mice. Exp Anim, 70, 145–160.

Menichella, D. M., Goodenough, D. A., Sirkowski, E., Scherer, S. S. & Paul, D. L. 2003. Connexins are critical for normal myelination in the Cns. J Neurosci, 23, 5963–73.

Mullen, R. J., Eicher, E. M. & Sidman, R. L. 1976. Purkinje cell degeneration, a new neurological mutation in the mouse. Proc Natl Acad Sci U S A, 73, 208–12.

Murakami, Y., Saito, K., Ito, H. & Hashimoto, Y. 2019. Transferrin isoforms in cerebrospinal fluid and their relation to neurological diseases. Proc Jpn Acad Ser B Phys Biol Sci, 95, 198–210.

Neusch, C., Rozengurt, N., Jacobs, R. E., Lester, H. A. & Kofuji, P. 2001. Kir4.1 potassium channel subunit is crucial for oligodendrocyte development and in vivo myelination. J Neurosci, 21, 5429–38.

Nishiyama, A., Shimizu, T., Sherafat, A. & Richardson, W. D. 2021. Life-long oligodendrocyte development and plasticity. Semin Cell Dev Biol, 116, 25–37.

Noveroske, J. K., Hardy, R., Dapper, J. D., Vogel, H. & Justice, M. J. 2005. A new ENU-induced allele of mouse quaking causes severe CNS dysmyelination. Mamm Genome, 16, 672–82.

Novodvorsky, P. & Chico, T. J. 2014. The role of the transcription factor KLF2 in vascular development and disease. Prog Mol Biol Transl Sci, 124, 155–88.

Rao, V. T. S., Khan, D., Cui, Q. L., Fuh, S. C., Hossain, S., Almazan, G., Multhaup, G., Healy, L. M., Kennedy, T. E. & Antel, J. P. 2017. Distinct age and differentiation-state dependent metabolic profiles of oligodendrocytes under optimal and stress conditions. PLos One, 12, e0182372.

Readhead, C. & Hood, L. 1990. The dysmyelinating mouse mutations shiverer (shi) and myelin deficient (shimld). Behav Genet, 20, 213–34.

Reinert, A., Morawski, M., Seeger, J., Arendt, T. & Reinert, T. 2019. Iron concentrations in neurons and glial cells with estimates on ferritin concentrations. BMC Neurosci, 20, 25.

Roach, A., Boylan, K., Horvath, S., Prusiner, S. B. & Hood, L. E. 1983. Characterization of cloned cdna representing rat myelin basic protein: absence of expression in brain of shiverer mutant mice. Cell, 34, 799–806.

Rosko, L. M., Gentile, T., Smith, V. N., Manavi, Z., Melchor, G. S., Hu, J., Shults, N. V., Albanese, C., Lee, Y., Rodriguez, O. & Huang, J. K. 2023. Cerebral Creatine Deficiency Affects the Timing of Oligodendrocyte Myelination. J Neurosci, 43, 1143–1153.

Saleh, M. C., Espinosa De Los Monteros, A., De Arriba Zerpa, G. A., Fontaine, I., Piaud, O., Djordjijevic, D., Baroukh, N., Garcia Otin, A. L., Ortiz, E., Lewis, S., Fiette, L., Santambrogio, P., Belzung, C., Connor, J. R., De Vellis, J., Pasquini, J. M., Zakin, M. M., Baron, B. & Guillou, F. 2003. Myelination and motor coordination are increased in transferrin transgenic mice. J Neurosci Res, 72, 587–94.

Schmidt, A., Marescau, B., Boehm, E. A., Renema, W. K., Peco, R., Das, A., Steinfeld, R., Chan, S., Wallis, J., Davidoff, M., Ullrich, K., Waldschütz, R., Heerschap, A., De Deyn, P. P., Neubauer, S. & Isbrandt, D. 2004. Severely altered guanidino compound levels, disturbed body weight homeostasis and impaired fertility in a mouse model of guanidinoacetate N-methyltransferase (GAMT) deficiency. Hum Mol Genet, 13, 905–21.

Sturrock, R. R. 1980. Myelination of the mouse corpus callosum. Neuropathol Appl Neurobiol, 6, 415–20.

Suzuki, K. & Zagoren, J. C. 1977. Quaking mouse: an ultrastructural study of the peripheral nerves. J Neurocytol, 6, 71–84.

Van De Wal, M. A. E., Adjobo-Hermans, M. J. W., Keijer, J., Schirris, T. J. J., Homberg, J. R., Wieckowski, M. R., Grefte, S., Van Schothorst, E. M., Van Karnebeek, C., Quintana, A. & Koopman, W. J. H. 2022. Ndufs4 knockout mouse models of Leigh syndrome: pathophysiology and intervention. Brain, 145, 45–63.

Yoshikawa, F., Sato, Y., Tohyama, K., Akagi, T., Furuse, T., Sadakata, T., Tanaka, M., Shinoda, Y., Hashikawa, T., Itohara, S., Sano, Y., Ghandour, M. S., Wakana, S. & Furuichi, T. 2016. Mammalian-Specific Central Myelin Protein Opalin Is Redundant for Normal Myelination: Structural and Behavioral Assessments. PLos One, 11, e0166732.

Yoshiki, A. & Moriwaki, K. 2006. Mouse phenome research: implications of genetic background. Ilar j, 47, 94–102.

Zeiss, C. J. 2021. Comparative Milestones in Rodent and Human Postnatal Central Nervous System Development. Toxicol Pathol, 49, 1368–1373.

Zhang, Y., Chen, K., Sloan, S. A., Bennett, M. L., Scholze, A. R., O’keeffe, S., Phatnani, H. P., Guarnieri, P., Caneda, C., Ruderisch, N., Deng, S., Liddelow, S. A., Zhang, C., Daneman, R., Maniatis, T., Barres, B. A. & Wu, J. Q. 2014. An RNA-sequencing transcriptome and splicing database of glia, neurons, and vascular cells of the cerebral cortex. J Neurosci, 34, 11929–47.

